# Evolutionary Implications of Anoxygenic Phototrophy in the Bacterial Phylum *Candidatus* Eremiobacterota (WPS-2)

**DOI:** 10.1101/534180

**Authors:** Lewis M. Ward, Tanai Cardona, Hannah Holland-Moritz

## Abstract

Genome-resolved environmental metagenomic sequencing has uncovered substantial previously unrecognized microbial diversity relevant for understanding the ecology and evolution of the biosphere, providing a more nuanced view of the distribution and ecological significance of traits including phototrophy across diverse niches. Recently, the capacity for bacteriochlorophyll-based anoxygenic photosynthesis has been found in the uncultured bacterial WPS-2 phylum (recently proposed as *Candidatus* Eremiobacterota) that are in close association with boreal moss. Here, we use phylogenomic analysis to investigate the diversity and evolution of phototrophic WPS-2. We demonstrate that phototrophic WPS-2 show significant genetic and metabolic divergence from other phototrophic and non-phototrophic lineages. The genomes of these organisms encode a completely new family of anoxygenic Type II photochemical reaction centers and other phototrophy-related proteins that are both phylogenetically and structurally distinct from those found in previously described phototrophs. We propose the name *Candidatus* Palusbacterales for the order-level aerobic WPS-2 clade which contains phototrophic lineages, from the Latin for “bog bacteria”, in reference to the typical habitat of phototrophic members of this clade.

## Introduction

The vast majority of primary productivity on Earth is fueled by photosynthesis, both today^1^ and through most of the history of life^2–4^, making the organisms and proteins responsible for driving photosynthesis into critical bases for the carbon cycle. While the quantitatively most significant form of photosynthesis today is oxygenic photosynthesis, which uses water as an electron donor to power carbon fixation, there is also a great diversity of bacteria capable of anoxygenic photosynthesis using compounds such as sulfide, molecular hydrogen, ferrous iron, or arsenic as electron donors^5,6^. Anoxygenic photosynthesis is restricted to members of the bacterial domain, where it has a scattered phylogenetic distribution (Figure 1). While the first observations of anoxygenic photosynthesis were made in the late 1800s^7^, the known diversity of bacteria capable of anoxygenic phototrophy has exploded in recent years thanks largely to genome-resolved metagenomics^8,9^ (Figure 1). As our understanding of the physiology and evolution of phototrophy is contingent on adequate sampling of the diversity of phototrophic organisms, continuing discoveries of phylogenetically, ecologically, and biochemically novel phototrophs spurs new insights into phototrophy today and in Earth’s history.

**Figure 1.**
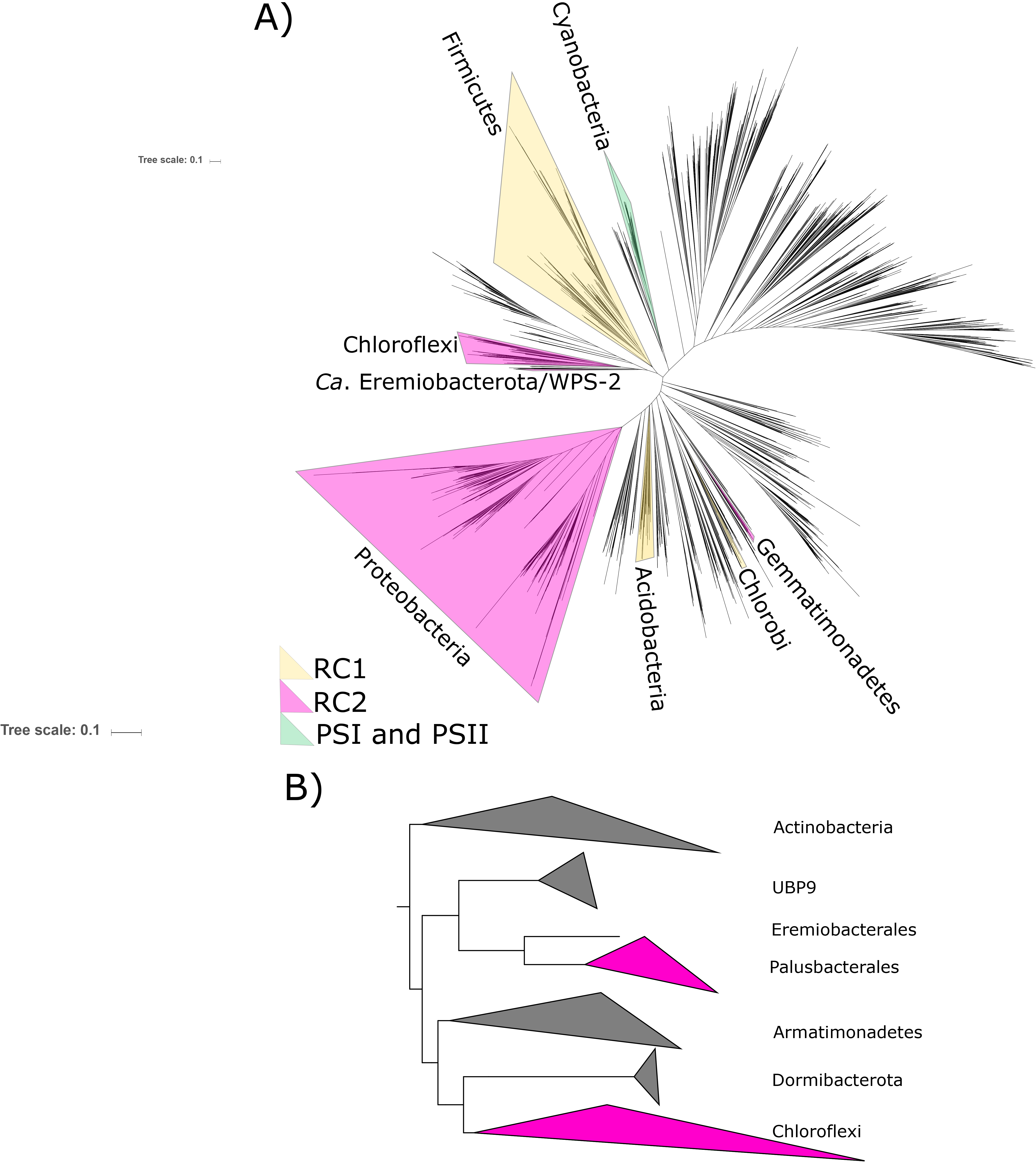
(A) Phylogenetic tree of bacteria showing position of WPS-2/*Ca*. Eremiobacterota, based on concatenated ribosomal protein sequences following Hug et al. 2016, with the distribution of photosynthetic reaction centers color-coded. In case where only some members of a phylum are capable of phototrophy (e.g. Heliobacteria in the much larger Firmicutes phylum), the entire phylum is highlighted for clarity. (B) Zoomed-in relationships of *Ca*. Eremiobacterota (here subdivided into candidate orders UBP9, Eremiobacterales, and Palusbacterales), Chloroflexi, and other closely related phyla.

Unlike oxygenic photosynthesis which requires the work of Type I and Type II photochemical reaction centers linked in series for water oxidation, anoxygenic phototrophs use exclusively either Type I or Type II reaction centers. Type I and II reaction centers can be differentiated by the nature of their electron acceptor^5^. Type I reduce ferredoxin, while Type II reduce quinones. There are only four phyla of bacteria known to have phototrophic representatives that use anoxygenic Type II reaction centers: Proteobacteria, Chloroflexi, Gemmatimonadetes^10^ and the newly discovered WPS-2^11^ (Figure 1). The photochemical pigments of anoxygenic Type II reaction centers are bound by two homologous subunits known as L (*PufL*) and M (*PufM*). These have traditionally been subdivided into two types, the L and M found in the reaction centers of phototrophic Proteobacteria (PbRC) and the L and M found in those of the phototrophic Chloroflexi (CfRC), with each set making separate monophyletic clades^12,13^. Therefore, substantial differences exist between the PbRC and the CfRC not only at the level of photochemistry, but also at the level of sequence identity of the core subunits, pigment and subunit composition^14,15^. Phototrophic Gemmatimonadetes, on the other hand, encode Proteobacteia-like reaction centers, as Zeng et al.^10^ demonstrated that this group obtained phototrophy via horizontal gene transfer of an entire photosynthesis gene cluster from a gammaproteobacterium.

Here we show that the genomes of the newly discovered phototrophs of the uncultivated candidate phylum WPS-2 (also known as *Candidatus* Eremiobacterota^16^) encode a third distinct lineage of anoxygenic Type II reaction centers with novel and unusual characteristics—only the third of this type after the discovery of the Chloroflexi nearly half a century ago^17^. We also show that the WPS-2 clade encoding phototrophy is distinct from basal Eremiobacterota and related phyla in terms of traits including the capacity for aerobic respiration, membrane architecture, and environmental distribution, reflecting a divergent evolutionary history and likely novel ecological roles.

## Methods

### Metagenome Analyses

Metagenome-assembled genomes (MAGs) of WPS-2 bacteria were downloaded from NCBI WGS and JGI IMG databases. Completeness and contamination of genomes was estimated based on presence and copy number of conserved single-copy proteins by CheckM^18^. Sequences of ribosomal and metabolic proteins used in analyses (see below) were identified locally with the *tblastn* function of BLAST+^19^, aligned with MUSCLE^20^, and manually curated in Jalview^21^. Positive BLAST hits were considered to be full length (e.g. >90% the shortest reference sequence from an isolate genome) with *e*-values greater than 1e^−20^. Genes of interest were screened against outlier (e.g. likely contaminant) contigs as determined by CheckM^18^ and RefineM^22^ using tetranucleotide, GC, and coding density content. Presence of metabolic pathways of interest was predicted with MetaPOAP^23^ to check for False Positives (contamination) or False Negatives (genes present in source genome but not recovered in metagenome-assembled genomes). Phylogenetic trees were calculated using RAxML^24^ on the Cipres science gateway^25^. Transfer bootstrap support values were calculated by BOOSTER^26^, and trees were visualized with the Interactive Tree of Life viewer^27^. Taxonomic assignment was confirmed with GTDB-Tk^28^ and by placement in a concatenated ribosomal protein phylogeny following methods from Hug et al.^29^ Histories of vertical versus horizontal inheritance of metabolic genes was inferred by comparison of organismal and metabolic protein phylogenies to determine topological congruence^9,30^.

### Reaction center evolution analyses

A total of 14 L and 12 M amino acid sequences were collected from the compiled metagenome data. The sequences were added to a dataset of Type II reaction center subunits compiled before^31^, which included sequences from Cyanobacteria, Proteobacteria, and Chloroflexi. Sequence alignments were done in Clustal Omega^32^ using 10 combined guide trees and Hidden Markov Model iterations. Maximum Likelihood phylogenetic analysis was performed with the PhyML online service (http://www.atgc-montpellier.fr/phyml/)^33^ using the Smart Model Selection and Bayesian information criterion for the computation of parameters from the dataset^34^. Tree search operations were performed with the Nearest Neighbor Interchange approach and the approximate likelihood-ratio test method was chosen for the computation of branch support values. A phylogenetic tree was also constructed using only L and M subunits and excluding cyanobacterial homologs.

Transmembrane helix prediction on WPS-2 L and M subunits was computed using the TMHMM Server, v. 2.0 (http://www.cbs.dtu.dk/services/TMHMM/)^35^ and the ΔG transmembrane helix prediction tool v. 1.0 (http://dgpred.cbr.su.se/)^36^. Structural homology models were carried out with SWISS-MODEL server (https://swissmodel.expasy.org/)^37^ using the 1.9 Å crystal structure of the gammaproteobacterium *Thermochromatium tepidum* as template^14^ (PDB ID: 5y5s). The WPS-2 L and M subunits annotated as Ga0175859_11240458 and Ga0175859_11402733 respectively were used for the structural reconstruction.

## Results and Discussion

### Ecology and distribution of WPS-2

Clonal libraries containing 16S rRNA sequences from the candidate WPS-2 (Wittenberg Polluted Soil) phylum were first identified in a 2001 study of soil polluted with polychlorinated biphenyl (PCB) in Germany^38^. Since then, WPS-2 has been found in 16S rRNA amplicon studies from diverse environments including acidic, polluted environments^38–40^, alpine, high latitude and antarctic soils^16,41–43^, cryoconite holes in Greenland^44^, human/canine oral microbiomes^45^, bogs and peatlands^46,47^. More recently, several metagenomic studies have successfully assembled MAGs from WPS-2. These studies include samples from an acidic mine, rich in arsenic^48^ (MAGs from this study were later identified as WPS-2 by Camanocha and Dewhirst^45^), Arctic^22^ and Antarctic desert soils^16^, boreal mosses^11^, and a peatland in Sweden^49^.

Although these environments are varied, several trends emerge: WPS-2 has a global distribution and is most often found in cool, acidic, and aerobic environments. While not exclusively found in low-temperature environments, most samples containing WPS-2 were collected from areas that are typically cold (i.e. Antarctica, Greenland), or undergo shorter growing seasons (high-latitude and alpine environments). Similarly, WPS-2 is more often found in sites with acidic to moderately acidic pH (between 3 and 6.5). In some cases, this acidity has come from pollution, in others, such as acidic springs, peatlands, bogs, and fir-spruce forest soil, it is a natural part of the habitat. Although WPS-2 has often been associated with polluted environments, it seems more likely that this is due the acidity accompanying these environments than to any special resistance to the varied pollutants from these studies. Oxygen-rich environments such as mosses, cryoconite holes, and the top few centimeters of soil are typical habitat for WPS-2 and several papers have speculated that members of this phylum are therefore aerobic or microaerobic^11,16,41^.

Often WPS-2 makes up a minor part of the bacterial community, however in some notable cases, the phylum has either dominated the bacterial community (e.g. 23-25%)^42-44^ or several phylotypes from WPS-2 have been among the dominant taxa^11,47^. Although most studies identifying WPS-2 as dominant are from specialized or unique environments (i.e. acidic springs, Antarctic soils, and cryoconite holes), an easily-accessible and commonplace environment in which phylotypes of WPS-2 are often abundant, is boreal mosses. Phylotypes from WPS-2 are the second-most dominant taxa across 6 common boreal moss species and absent in only one moss of the 7 species studied^11^. Interestingly, even when mosses are not the focus of a study, WPS-2 is often more common in environments that contain them^40,46,47,49^. Currently there are no isolates of WPS-2 to investigate its biochemistry, or to use in comparative genomics, yet given its phototrophic properties and evolutionary context, such an isolate would be useful. As an accessible and abundant host to phylotypes of WPS-2, boreal mosses provide a logical starting place from which to try to culture WPS-2—an essential step in upgrading this candidate phylum to taxonomically valid status as well as for characterizing its physiology.

### Phylogenetic context of phototrophy in the WPS-2 phylum

The WPS-2 phylum is located on a branch of the bacterial tree of life located near the phyla Chloroflexi, Armatimonadetes, and Dormibacteria (Figure 1). This placement has interesting implications for the evolutionary history of a variety of traits (as discussed below) but is particularly relevant for considerations of the evolutionary history of anoxygenic phototrophy.

Phylogenetic analysis of proteins for the Type II reaction center suggests that phototrophy in WPS-2 might be most closely related to that of the Chloroflexi (Figure 2, but see below). Given the relatedness of WPS-2 and Chloroflexi phototrophy proteins (Figure 2, Supplemental Figure S1) and their relative closeness in organismal phylogenies (Figure 1), an important outstanding question is whether the last common ancestor of these phyla was phototrophic, implying extensive loss of phototrophy in both phyla and their relatives, or whether WPS-2 and Chloroflexi have independently acquired phototrophy via HGT subsequent to the divergence of these lineages (likely from an earlier branching uncharacterized or extinct group of phototrophic bacteria). While this cannot be conclusively ascertained with available data, it is apparent that aside from bacteriochlorophyll synthesis and phototrophic reaction centers, necessary proteins involved in phototrophic electron transfer (e.g. a *bc* complex or alternative complex III) are not shared, suggesting that members of these phyla independently acquired respiratory electron transfer necessary for phototrophy after their divergence. This is consistent with a relatively recent origin of phototrophy in the Chloroflexi (i.e. within the last ~1 billion years) after the acquisition of aerobic respiration by the Chloroflexi and other phyla following the Great Oxidation Event (GOE) ~2.3 billion years ago^50,51^. Together with the relatively high protein sequence similarity between reaction center proteins among the phototrophic WPS-2 (discussed below) this may suggest that phototrophy in the WPS-2 has radiated on a similar or even shorter timescale.

**Figure 2.**
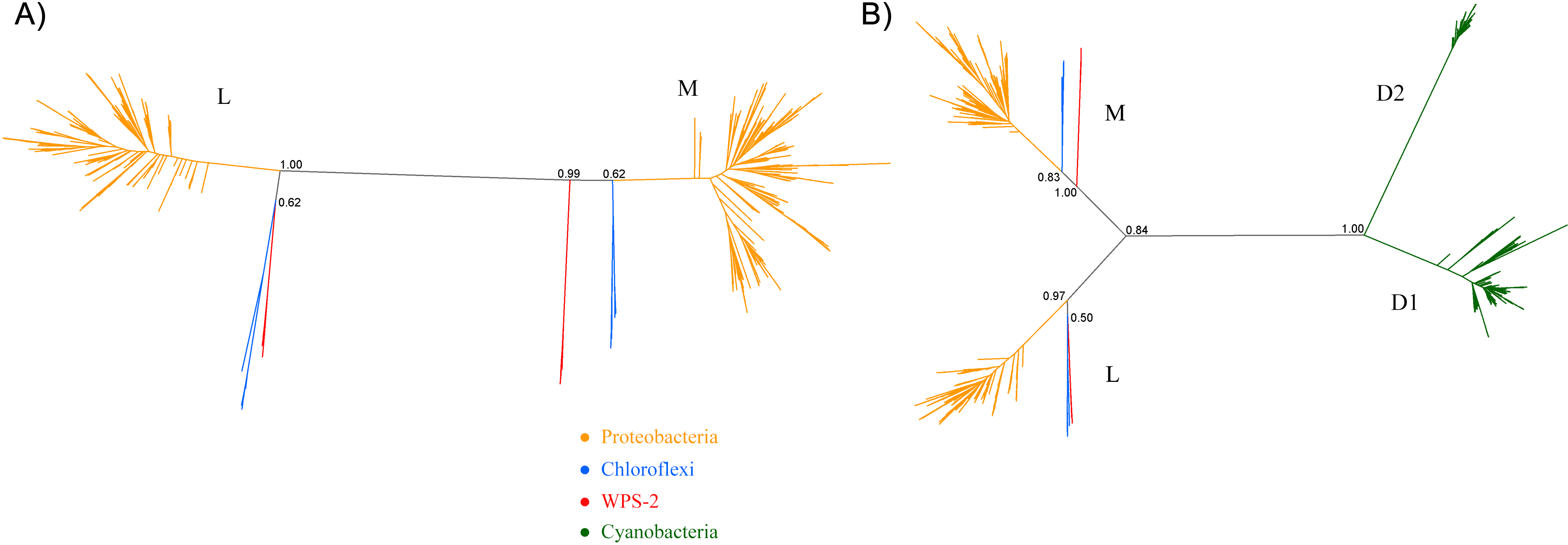
Phylogeny of Type II reaction center proteins. (A) A Tree calculated using only L and M subunits including only anoxygenic phototrophs. (B) A Tree including D1 and D2 proteins of Cyanobacteria.

Phototrophy in WPS-2 is not restricted to a single lineage, but appears in at least four discrete lineages interspersed with nonphototrophic lineages within the phylum (Figure 3). The topology of the organismal phylogeny of the WPS-2 is incongruent with that of phototrophy protein phylogenies (Figure 3, Supplemental Figure S1), and so this distribution may best be explained by a history of HGT between members of the WPS-2 and not by a deeper ancestry of phototrophy followed by extensive loss. Similar cases of multiple HGT events of complete photosynthesis gene clusters have been demonstrated within the family Rhodobacteraceae of Alphaproteobacteria^52^, between classes of the phylum Chloroflexi^9^, and into the Gemmatimonadetes phylum from members of the Proteobacteria^10^, which might suggest convergent evolutionary patterns driving the distribution of Type II reaction center-based anoxygenic phototrophy within distant lineages.

**Figure 3.**
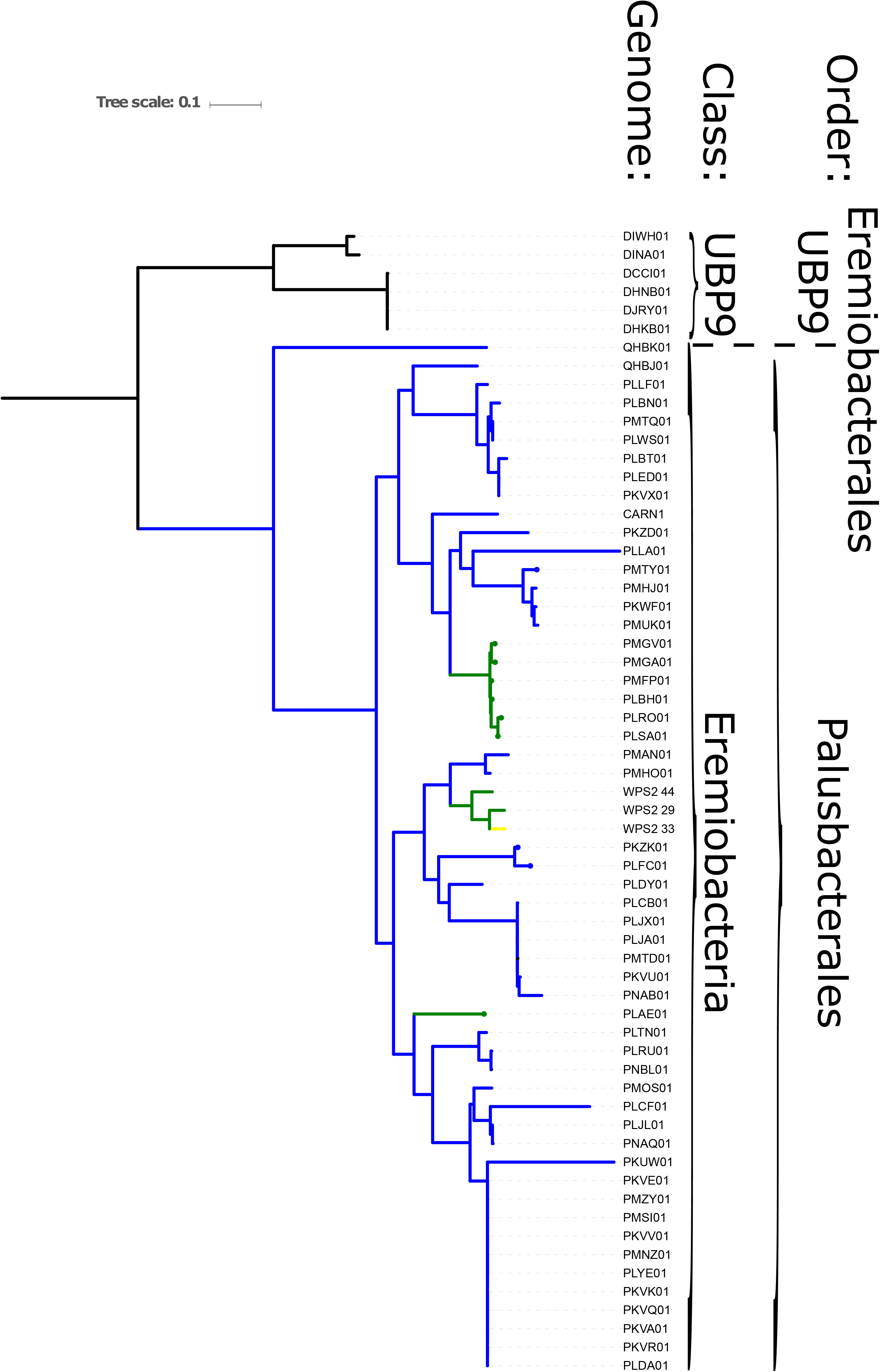
Phylogeny of WPS-2 built with concatenated ribosomal protein sequences, annotated with distribution of traits including phototrophy and aerobic respiration. Green branches encode both phototrophy and aerobic respiration; blue lineages encode aerobic respiration but not phototrophy; yellow lineages encode phototrophy but not aerobic respiration; black lineages encode neither.

The phylogenetic position of bacteriochlorophyll synthesis proteins including BchL of WPS-2 (Supplemental Figure S2) is somewhat in agreement with that of reaction center proteins (Figure 2, Supplemental Figure S1). However, BchL from the phylum Chlorobi is more closely related to that of the Chloroflexi than either is to the WPS-2. This is consistent with previous studies showing that HGT of protochlorophyllide and chlorophyllide reductase between stem group phototrophic Chlorobi and Chloroflexi had occurred^53,54^, although the direction of transfer had remained ambiguous. The new topology of BchL including WPS-2 suggests exchange of bacteriochlorophyll synthesis proteins via HGT from stem group phototrophic Chloroflexi to stem group phototrophic Chlorobi. It is likely that exchange of photosynthetic components between Chloroflexi and Chlorobi had occurred more than once, in either direction, and at different time points and may have involved a swap of reaction center type as suggested before^53^ and the acquisition of chlorosomes^55,56^.

### Additional metabolic traits in WPS-2 genomes

Aside from genes encoding phototrophy, the phototroph-containing WPS-2 clade is distinct from its relatives due to a variety of metabolic traits, including proteins used for aerobic respiration and carbon fixation, as well as evidence for an outer membrane.

Most WPS-2 genomes encode aerobic respiration via an A-family heme-copper oxidoreductase (HCO) and a *bc* complex III. This is in contrast to the Chloroflexi, which typically encode an alternative complex III instead of a *bc* complex. Protein sequences of respiratory proteins from WPS-2 genomes primarily form single closely related clades with topologies that broadly reflect organismal relationships (Figure 3, Supplemental Figures S3 and S4); these trends suggest vertical inheritance of these genes from the last common ancestor of this clade. The main exceptions to these trends are the divergent WPS-2 genomes in the UNP9 class, which appear to be ancestrally anaerobic, and a handful of WPS-2 genomes that encode additional copies of respiratory proteins, which appear to have been acquired via horizontal gene transfer more recently and have since undergone additional HGT between members of the WPS-2.

Like the Chloroflexi, phototrophic WPS-2 typically encode a B-family HCO in addition to the ancestral A-family HCO used for respiration at relatively high O_2_ concentrations^9^; the functional relationship between the B-family O_2_ reductase and phototrophy is unclear, but may relate to oxygen sensitivity of phototrophy-related proteins and the greater efficacy of B-family HCOs at low oxygen concentrations^57^.

While phototrophic Chloroflexi typically perform carbon fixation via the 3-hydroxypropionate bicycle^50^, carbon fixation in phototrophic WPS-2 is encoded via a Form I rubisco. All phototrophic genomes in this phylum encode closely related rubisco proteins related to that of *Kouleothrix aurantiaca*^9^, with some also encoding a second Form I rubisco copy more closely related to sequences from *Bradyrhizobium* (Supplemental Figure S5). The PMHO01 and PMHP01 genomes encode rubisco despite not being phototrophic; this rubisco is not closely related to those encoded by phototrophic WPS-2, and so may reflect the independent acquisition of carbon fixation in some nonphototrophic WPS-2 lineages (perhaps to support a lithoautotrophic lifestyle). The phylogeny of rubisco proteins is not congruent with organismal or phototrophy trees, suggesting that carbon fixation may have undergone an independent history of HGT, consistent with trends in the Chloroflexi^9,50^.

The Chloroflexi are thought to possess a modified monoderm membrane^58,59^. WPS-2 genomes recovered genes for lipopolysaccharide synthesis and outer membrane proteins such as BamA, indicating that these organisms are diderm (i.e. possess an outer membrane), reinforcing interpretations of an ancestral diderm membrane architecture followed by secondary loss of the outer membrane in the phylum Chloroflexi^9,60–62^.

### Type II reaction centers in WPS-2

The phylogeny of Type II reaction center proteins showed that the L and M subunits of WPS-2 are distant, and clearly distinct, to those found in Proteobacteria and Chloroflexi (Figure 2), but the topology of the tree was unstable. It showed varying positions and levels of support for the WPS-2 sequences (Supplemental Figure S6). However, the sequence and structural comparison do reveal a slightly greater affinity and similarity to those of the Chloroflexi. The reaction center subunits show considerable diversity, with the level of sequence identity between the two most distant L and two most distant M subunits being about 59%. In comparison, the level of sequence identity between L or M subunits between two relatively distant strains of Chloroflexi, *Roseiflexus* spp. and *Chloroflexus* spp., is about 45% sequence identity. If the obtained sequences are representative of the diversity of phototrophic WPS-2, the greater sequence identity could potentially indicate a relatively more recent origin for their common ancestor in comparison to phototrophic Chloroflexi.

In addition to L and M subunits, the assembled genomes of WPS-2 also contain *pufC*, *puf2A* and *puf2C* encoding the tetraheme cytochrome direct electron donor to the photochemical pigments, and the light harvesting complex alpha and beta subunits, respectively^11^. No *puhA* was observed indicating that the reaction center of WPS-2 might lack an H subunit in a manner similar to the CfRC.

Sequence and structure prediction of the L and M subunits revealed a large number of unique characteristics (Figure 4), which is consistent with the level of distinctness seen in the phylogenetic analysis. In Proteobacteria and Cyanobacteria each Type II reaction center protein is made of 5 transmembrane helices each. The recent Cryo-EM structure of the reaction center from *Roseiflexus castenholzii* resolved a 6^th^ helix in the L subunit^15^. This was located at the N-terminus and cannot be detected through secondary structure prediction methods. In the case of WPS-2 L subunit, transmembrane helix prediction only detected the standard 5 helices; instead a novel N-terminal 6^th^ helix was detected in the M subunit (Figure 4 A).

**Figure 4.**
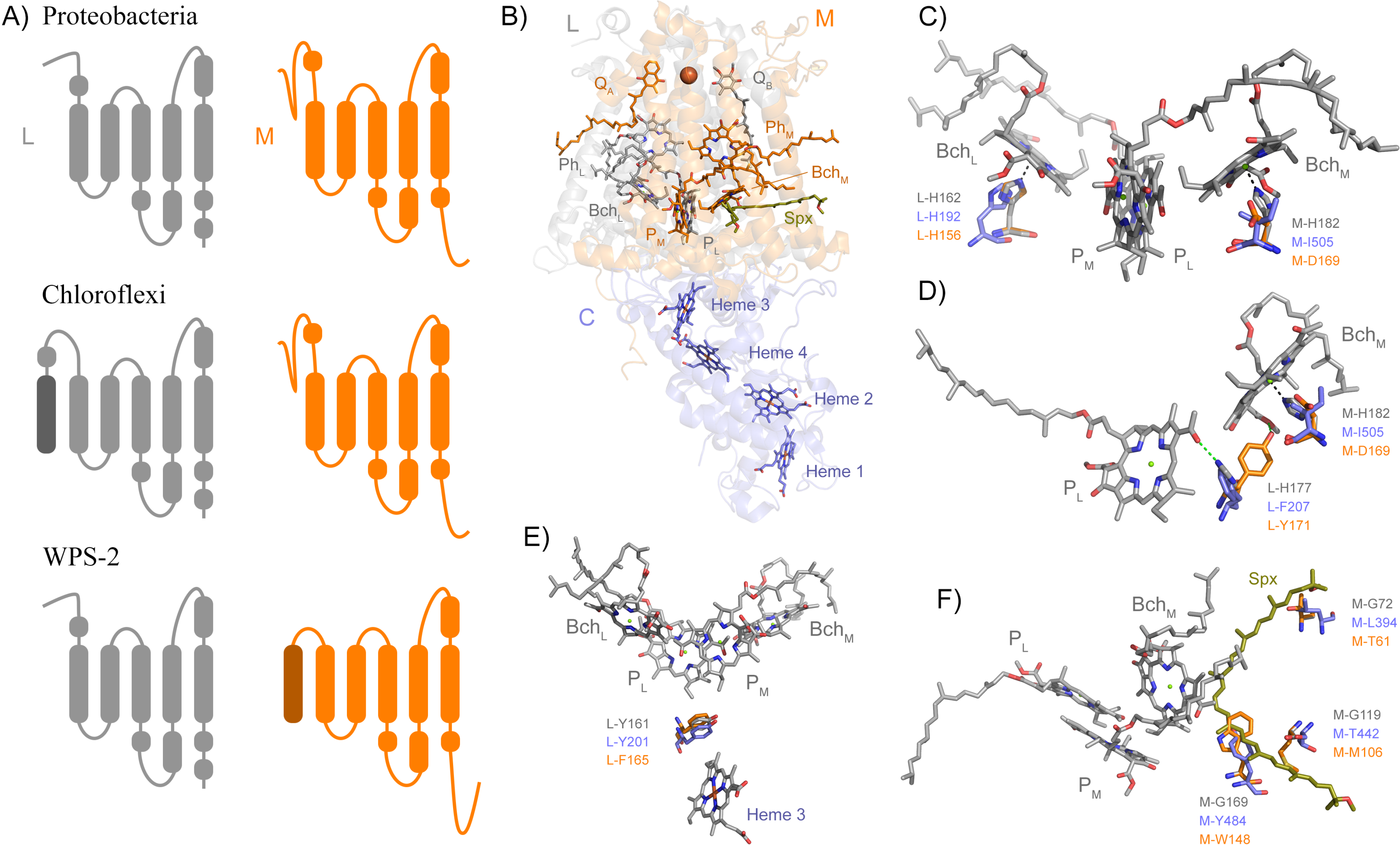
Structural comparisons of Type II reaction centers. (A) Schematic representation of the transmembrane helices. (B) The crystal structure of the PbRC from *Thermocromatium tepidum* highlighting all redox cofactors. (C) Comparison in the ligands to Bch_L_ and Bch_M_. Residues in grey are those from the PbRC, in blue those from the Cryo-EM structure of *Roseiflexus*, and in orange those from the homology model of the WPS-2 reaction center proteins. (D) Changes around hydrogen-bonding partners to P_L_. (E) Changes around L-Y161, a residue important for electron transfer from the tetraheme cytochrome into the oxidized photochemical pigments.

Most of the ligands to the photochemical pigments and cofactor are conserved, but unique changes that might modulate the energetics of electron transfer can be noted. For example, in the PbRC both “monomeric” bacteriochlorophylls, Bch_L_ and Bch_M_, are coordinated by histidine ligands (Figure 4 B and C). In the CfRC, a “third” bacteriopheophytin^15,63^ occupies the position of Bch_M_ and accordingly there is no histidine ligand, and instead an isoleucine is found. In WPS-2 a histidine ligand to Bch_M_ is also absent like in the CfRC, however instead of isoleucine and strictly conserved aspartate (M-D169) is found, which may suggests the presence of bacteriochlorophyll rather than bacteriopheophytin at this position, but with modified energetics^64^. This M-D169 might play an important role in the inactivation of electron transfer via the M branch.

In the PbRC, L-H177 provides a hydrogen bond to the photochemical pigment P_L_ (Figure 4 D). Mutagenesis studies have shown that the strength or absence of this bond modulates the midpoint potential of the primary donor pigments P_L_ and P_M_^65^. In the CfRC, the histidine is substituted by phenylalanine, which should contribute to the lower oxidizing potential of P in comparison to the PbRC. In the CfRC the midpoint potential of the primary donor is +360-390 mV^66–68^ while in the PbRC is +450-500 mV^69–72^. In the WPS-2 reaction center, a tyrosine is found at this position and it is potentially oriented towards Bch_M_ effectively breaking the hydrogen-bond to P_L_ like in the CfRC. The absence of the hydrogen-bond to P_L_ could lower the midpoint potential of the primary donor by about −95 mV^65^, which may suggests that WPS-2 has a primary donor with a potential more similar to that of the CfRC.

In the PbRC, a carotenoid is found in contact with Bch_M_ and it is thought that this protects the system against the formation of reactive oxygen species by quenching bacteriochlorophyll triplets^73^, see Figure 4 F. In the CfRC, no carotenoid has been reported at this position and the structure from *Roseiflexus* did not show a bound carotenoid^15^. In the CfRC and the WPS-2 reaction center three glycine residues, which in the PbRC give space to the carotenoid, are replaced by bulkier residues that should hinder the binding of a potential carotenoid.

In the PbRC and the CfRC a strictly conserved tyrosine (L-Y162) is located in between heme 3 (c-559) of PufC and the special pair (Fig. 5 E). Mutagenesis studies have shown that this tyrosine modulates the midpoint potential of both the heme and the special pair, but it is not required for fast electron transfer^74^. While a gene was identified for PufC in WPS-2^11^, this tyrosine was found to be a phenylalanine in all WPS-2 L sequences. A deprotonation of this conserved tyrosine residue stabilizes the primary charge separation reactions^75^, therefore this mechanism is not operational in the reaction center of WPS-2.

## Conclusions

Bacterial genomes previously assigned to WPS-2 have been assigned to a candidate phylum named *Candidatus* Eremiobacterota^16^. Recent genome-resolved metagenomics has expanded the known genetic diversity of this phylum^49^ as well as revealing the capacity for anoxygenic phototrophy^11^. Based on the metabolic traits of various *Ca*. Eremiobacterota lineages as discussed above, together with analysis via GTDB-Tk^28^ (Supplemental Table S1), we propose for the aerobic WPS-2 clade that includes phototrophic members the designation of a candidate order “Palusbacterales” (from the Latin for “bog”, in reference to the typical habitat of these organisms) within the Eremiobacteria class of Ca. Eremiobacterota. Following current standards for the taxonomy of uncultured taxa^76^, we propose the genome WPS2_44 (IMG accession #2734482170) as the type species *Palusbacter phototrophicum* (from the Latin for “light-eating bog bacterium”) for the candidate ranks Palusbacterales (order) and Palusbacteraceae (family).

Most lineages of anoxygenic phototrophs can be preferentially found in characteristic environments, such as anoxic regions of stratified water columns for Chlorobi^77^, low-oxygen hot springs for Chloracidobacteria and phototrophic Chloroflexi^8,9,78–80^, and soils for Heliobacteria^81^, though exceptions do occur (e.g. Chloroflexi in carbonate tidal flats^82^ and Heliobacteria in hot springs^83^ or soda lakes^84,85^). In contrast, the phototrophic Proteobacteria and Gemmatimonadetes appear to have a more cosmopolitan distribution, including freshwater, marine, and soil environments^86,87^. The preferred environment for phototrophic members of *Ca.* Eremiobacterota appears to be cold, acidic, and aerobic environments with at least some members of the group forming a close association with plants, particularly mosses. This niche overlaps with that of plant-associated phototrophic Proteobacteria^88^, and may be due to the relatively high oxygen tolerance of these phototrophic lineages as compared to typically more oxygen-sensitive phototrophs such as Heliobacteria and Chlorobi. The apparent close and specific association of phototrophic *Ca.* Eremiobacterota with plants may reflect a long-term evolutionary association, which could further suggest that this group has radiated alongside plants over a timescale of <0.5 billion years, though this hypothesis will require molecular clocks or other analyses to test.

Most members of the class *Ca.* Eremiobacteria (i.e. the candidate orders Palusbacterales and Eremiobacterales) encode aerobic respiration using closely related A-family heme-copper oxidoreductases and *bc* complex III proteins (~85% of all genomes, and ~97% of >80% completeness) (Supplemental Figures S3 and S4). This is in contrast to the basal UBP9 class which appears to be composed of obligate anaerobes, with no respiration genes encoded in any of the available genomes in this clade, and aerobic Chloroflexi which do not encode closely related complex III or complex IV proteins. This, together with the broad congruence of *Ca.* Eremiobacteria organismal phylogenies with complex III and complex IV protein phylogenies, implies that aerobic respiration was acquired via horizontal gene transfer into stem group *Ca.* Eremiobacteria after their divergence from UBP9 but before the radiation of crown group Eremiobacteria. It is likely that anaerobic ancestors of the Eremiobacteria and UBP9 classes diverged diverged during Archean time, with the radiation of crown group *Ca.* Eremiobacteria occurring after the GOE ~2.3 billion years ago which led to the expansion of aerobic metabolisms^51^. This hypothesis will require molecular clock estimates for the divergence and radiation of *Ca.* Eremiobacteria or other tests to verify, but similar evolutionary trends have been seen in other groups, including the Chloroflexi and Cyanobacteria^50,51,89^. Following the acquisition of aerobic respiration by the last common ancestor of crown group *Ca.* Eremiobacteria, aerobic respiration via an A-family HCO and a *bc* complex III appears to have been largely vertically inherited, with additional respiratory proteins (e.g. second copies of *bc* complex and A-family HCOs, and the B-family HCOs associated with phototrophy) acquired via later HGT.

It is well established that all reaction centers originated as homodimers, with electron transfer occurring symmetrically on both side of the reaction center as it is still seen in Type I reaction centers^90^. It is thought that heterodimeric Type II reaction centers evolved from two independent duplication events, one leading to L and M in anoxygenic Type II and another leading to D1 and D2 in cyanobacterial PSII^12,13,91^. The heterodimerization process led to the evolution of asymmetric electron transfer, which occurs exclusively via the L branch in anoxygenic Type II and via D1 in PSII, while the M and D2 branches became inactive, respectively. That the three different anoxygenic Type II reaction centers show distinct mechanisms of redox tuning on key positions around Bch_M_ and other photochemical pigments could indicate that these lineages radiated soon after the L and M duplication. Furthermore, given that anoxygenic Type II reaction centers show up to five times faster rates of evolution than cyanobacterial PSII^31^ and that the emergence of at least some of major clades of phototrophs with anoxygenic Type II reaction centers postdate the GEO^50^, it is possible that the duplication of L and M subunits, and the radiation of the known anoxygenic Type II reaction centers occurred after the origin of oxygenic photosynthesis^31^. Therefore, the idea that extant forms of anoxygenic phototrophy powered by heterodimeric Type II reaction centers represent a “primitive” form of photosynthesis is not supported by the available data. Instead, it should be considered a relatively recent, highly specialized, and mobile evolutionary innovation within the larger context of the evolution of anoxygenic photosynthesis, which reaches into Paleoarchean time^92,93^. If so it would imply that anoxygenic photosynthesis early in Earth’s history may have been driven by now-extinct groups of bacteria using variations on phototropic pathways not yet observed today^94^.

The known phylogenetic diversity of anoxygenic phototrophs has increased substantially in recent years, thanks largely to genome-resolved metagenomic sequencing of diverse environments^8,9,11^. Better coverage of the extant diversity of phototrophs and their relatives allows us to better query the evolutionary history of both organisms and their metabolisms by providing opportunities for comparative genomic, phylogenetic, and molecular clock analyses. Previous attempts to interpret the early evolution of photosynthesis relied on the narrower set of phototrophic lineages that were known at the time (e.g.,^95,96^, whose analysis predated the discovery of phototrophic Chloracidobacteria, Gemmatimonadetes, Thermofonsia, and Eremiobacterota). The discovery of phototrophic Eremiobacterota and other lineages will provide crucial data for future reconsideration of the evolutionary relationships and history of phototrophy, improving our understanding of one of Earth’s most important metabolic pathways.

## Author’s contributions

L.M.W., T.C. and H.H.-M. designed the study, analyzed the data, and wrote the manuscript.

## Acknowledgements

T.C. is supported by a Leverhulme Trust grant (RPG-2017-223). L.M.W. is supported by an Agouron Institute postdoctoral fellowship. H. H. -M. is supported by NSF Dimensions of Biodiversity grant (DEB 1542609).

## Supplementary figures

**Figure S1.** Protein phylogeny of PufL and PufM subunits of the Type II reaction center of WPS-2 bacteria. Protein sequences that were incomplete due to metagenome assembly (<80% of full length) omitted.

**Figure S2.** Protein phylogeny of BchL proteins from phototrophic bacteria. Phyla labeled, and WPS-2 highlighted in red. Protein sequences that were incomplete due to metagenome assembly (<80% of full length) omitted.

**Figure S3.** Protein phylogeny of A- and B-family heme-copper oxidoreductases from WPS-2, Chloroflexi, and other related phyla. WPS-2 highlighted in red. Protein sequences that were incomplete due to metagenome assembly (<80% of full length) omitted.

**Figure S4.** Protein phylogeny of *bc* complex III from WPS-2, Chloroflexi, and other related phyla. WPS-2 highlighted in red. Protein sequences that were incomplete due to metagenome assembly (<80% of full length) omitted.

**Figure S5.** Protein phylogeny of the large subunit of rubisco from WPS-2, Chloroflexi, and other related phyla. Major clades outside of the WPS-2 collapsed for clarity. Protein sequences that were incomplete due to metagenome assembly (<80% of full length) omitted.

**Figure S6.** Protein phylogeny of reaction center proteins on a subset of 176 L and M sequences (A) and the same dataset but removing poorly aligned regions (B). Poorly aligned regions were removed with Gblocks allowing for smaller final blocks, gap positions within the final blocks, and less strict flanking positions.

### Supplemental tables

**Table S1.** Table of available WPS-2 genomes, including completeness, contamination, presence of respiration and phototrophy genes, candidate phylum assignment, metagenome source, and reference information.

### Supplemental Discussion

MetaPOAP False Negative estimates for the probability of phototrophy in each WPS-2 genome is found in Table 1; genomes in which some phototrophy proteins were recovered have a high probability of encoding the complete phototrophy pathway, whereas the probability that phototrophy is encoded in other genomes but was not recovered is very low, suggesting that the distribution of phototrophy shown in Figure 3 reflects the actual distribution of phototrophy in this phylum. Only the PKZK01 genome has a low (<0.01) MetaPOAP False Negative probability of encoding a full phototrophy pathway due to encoding only a subset of necessary proteins for phototrophy despite relatively high completeness; this organism may be an instance of transitional loss of phototrophy, but this will require more complete genome sequencing or physiological data to verify. While some Eremiobacteria genomes did not recover O_2_ reductases (PMTD01, PMUE01, PNBL01, and WPS2_33), these were primarily lower completeness (<80%), were closely related to aerobic strains, and did recover a *bc* complex, suggesting that these genomes either encode a respiratory electron transport chain that was not recovered in the MAG or that they are undergoing secondary loss of an ancestral respiratory metabolism.

