## Supplementary figures and images for "Evolutionary Implications of Anoxygenic Phototrophy in the Bacterial Phylum *Candidatus* Eremiobacterota (WPS-2)"

### Supplemental Figure S1

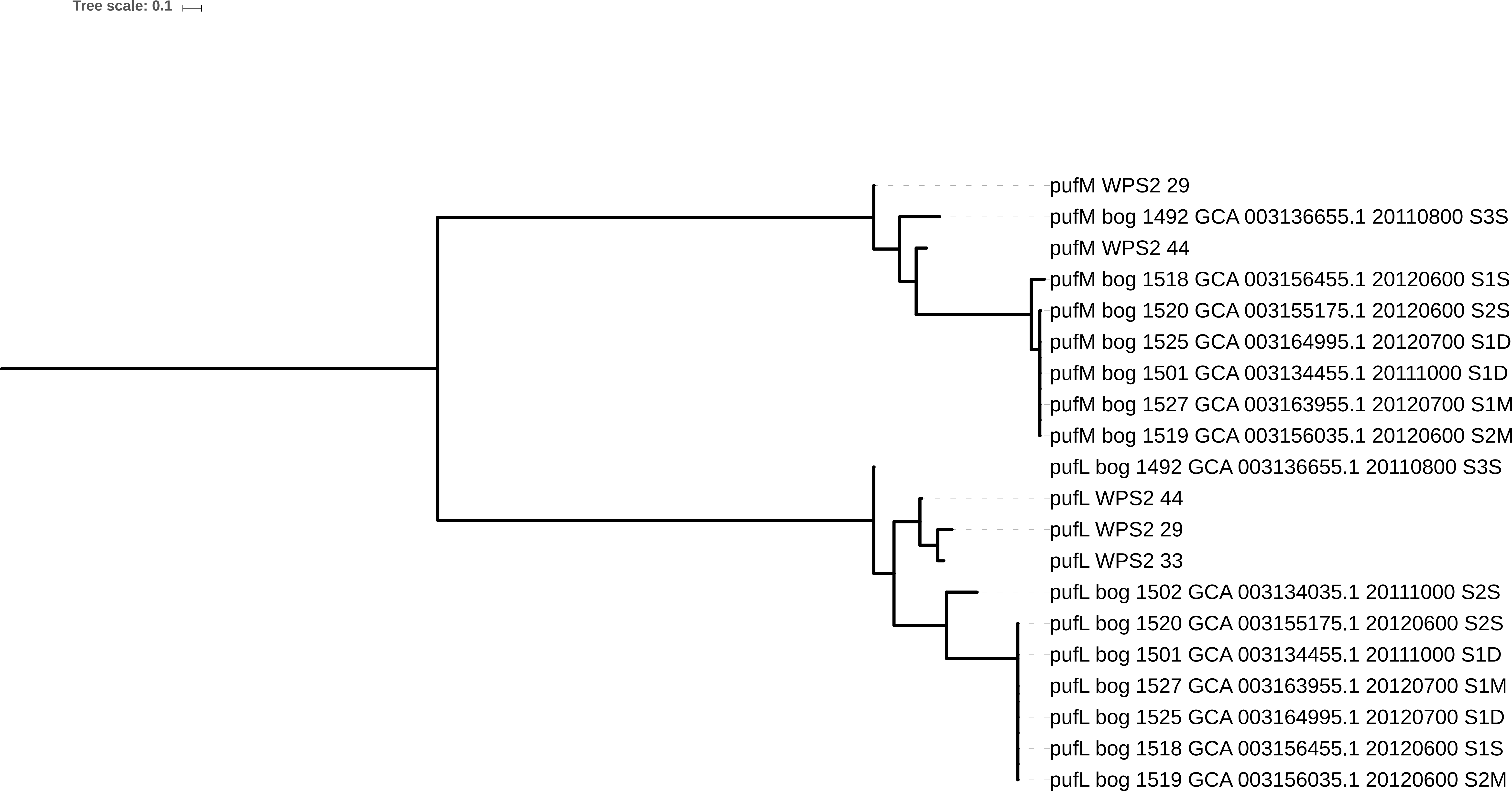

### Supplemental Figure S2

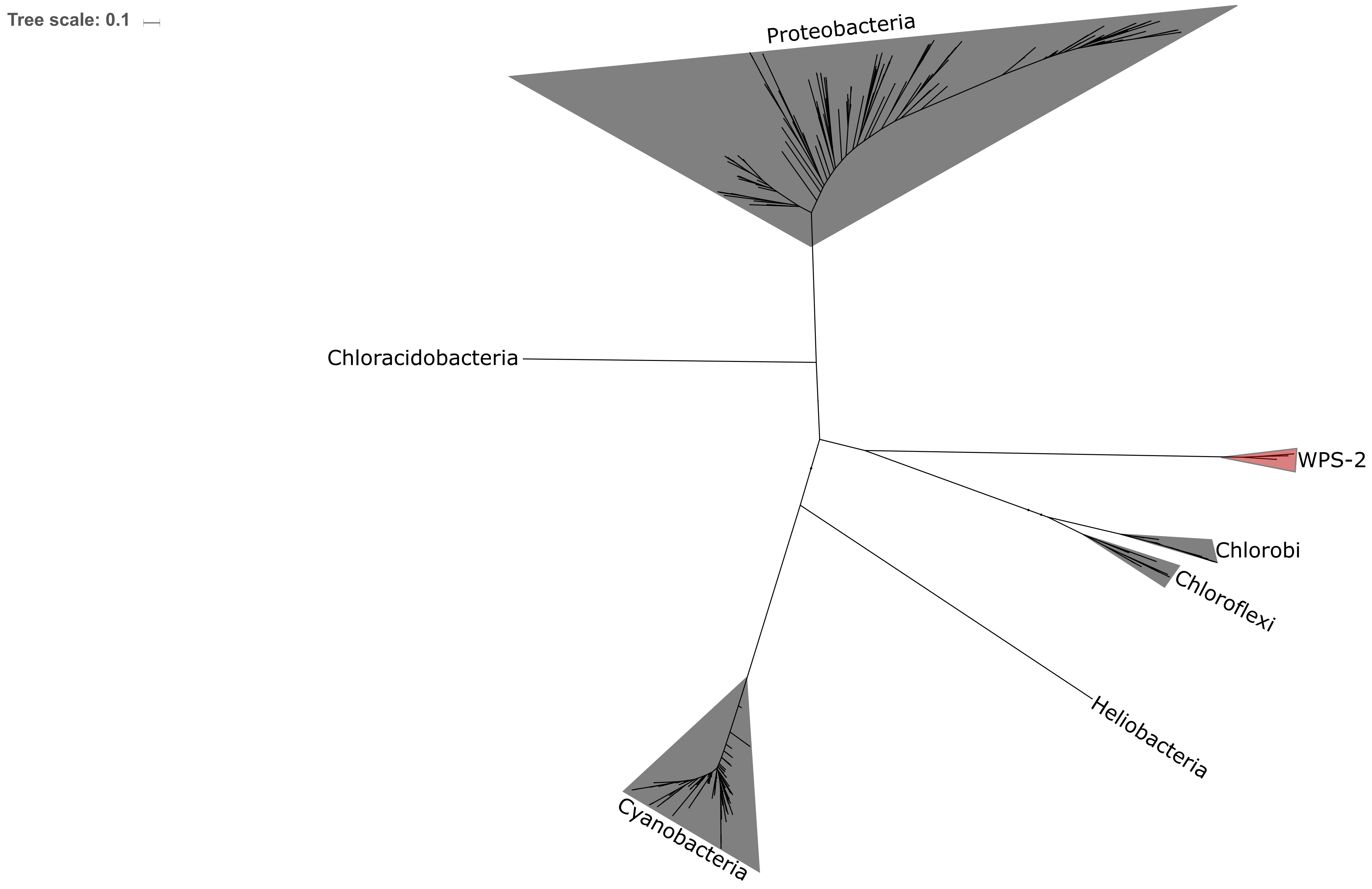

### Supplemental Figure S3

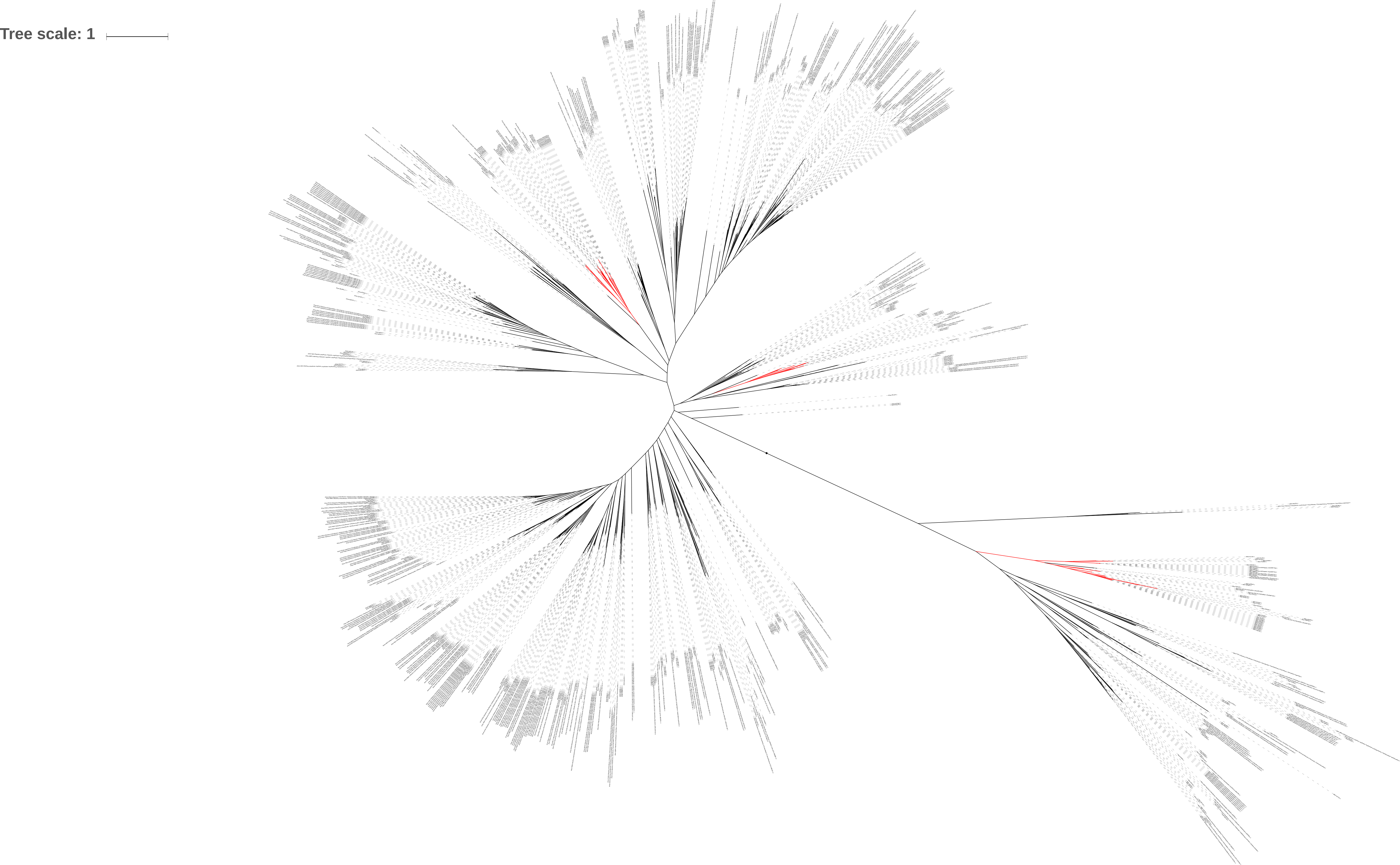

### Supplemental Figure S4

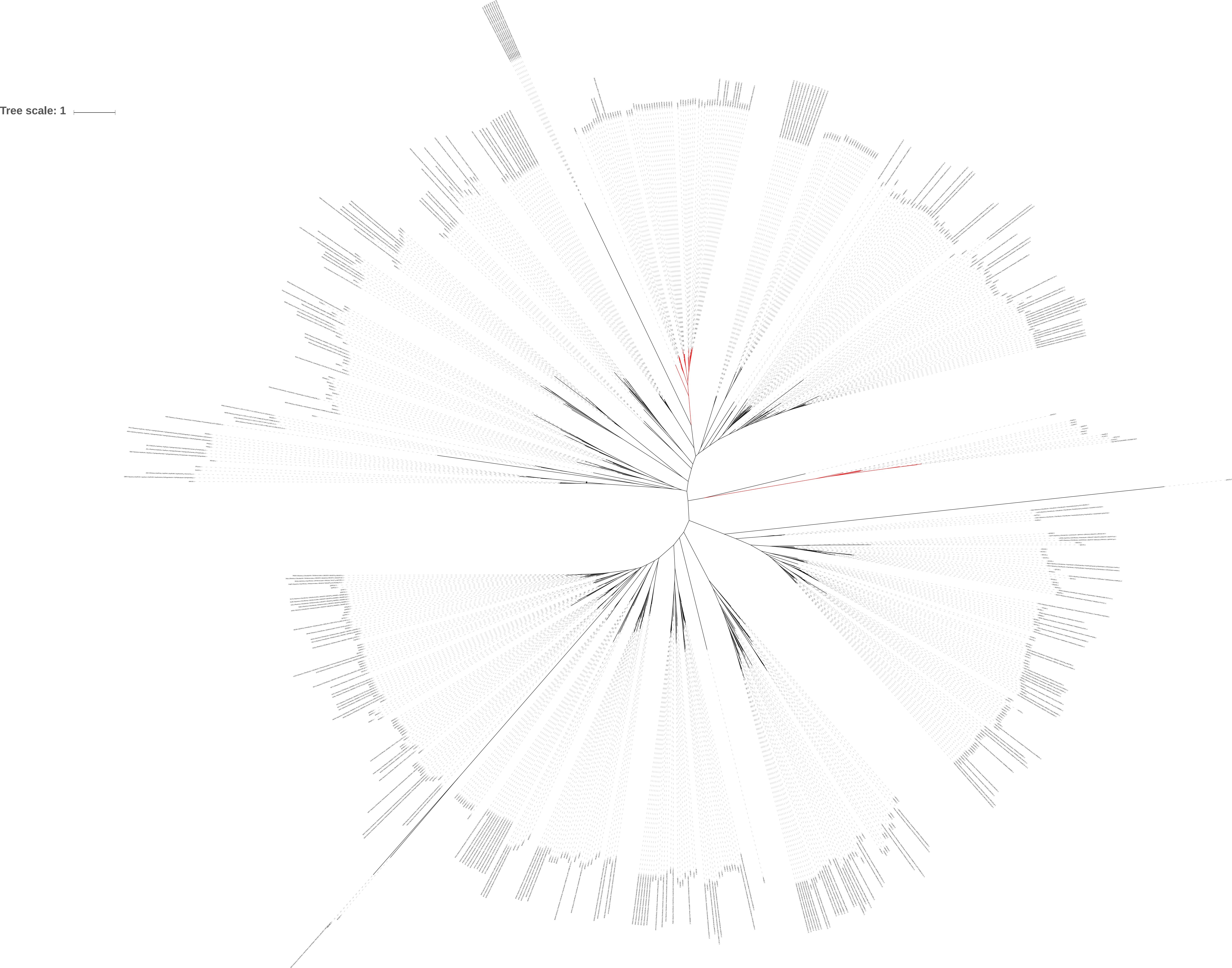

### Supplemental Figure S5

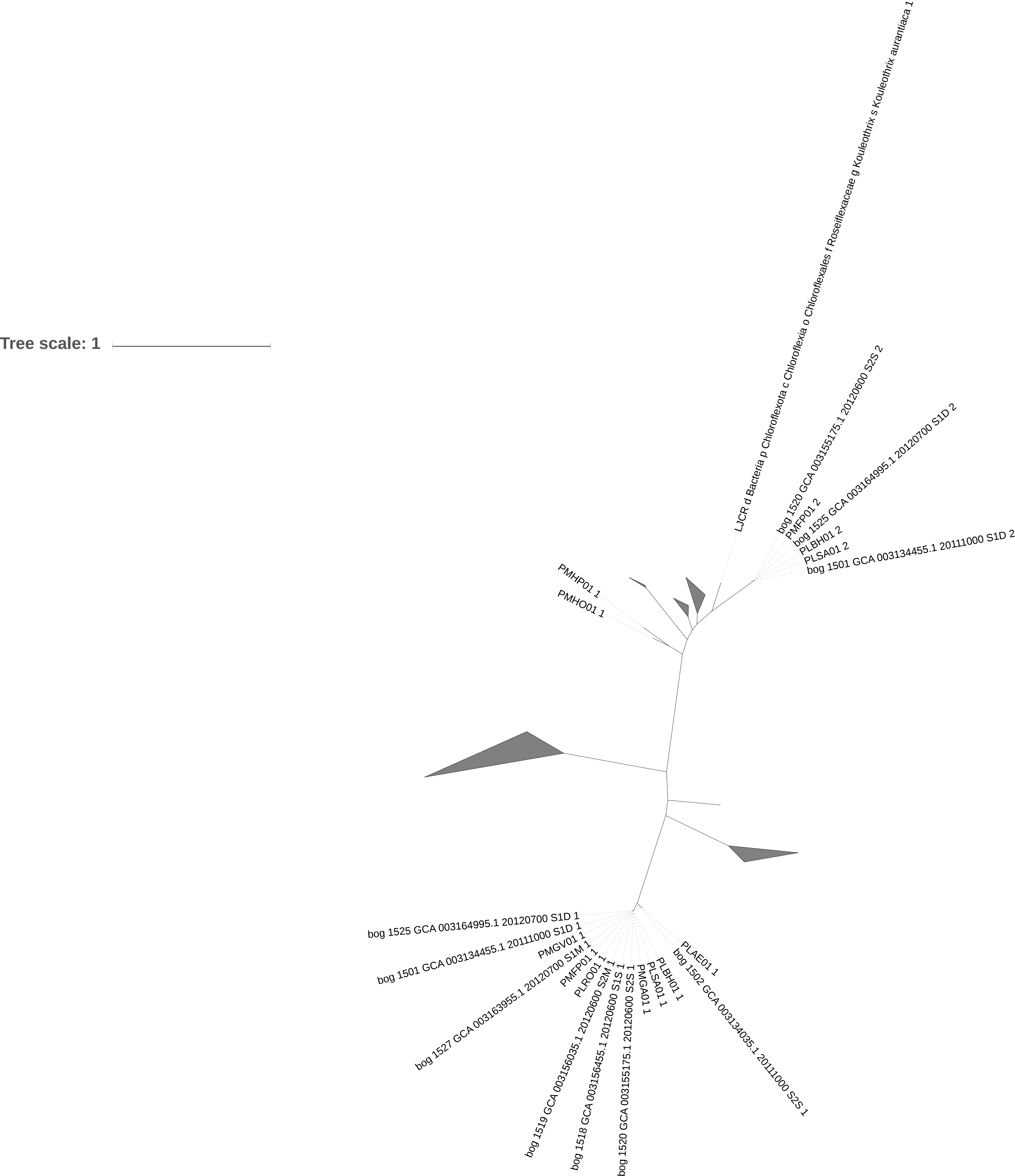

### Supplemental Figure S6

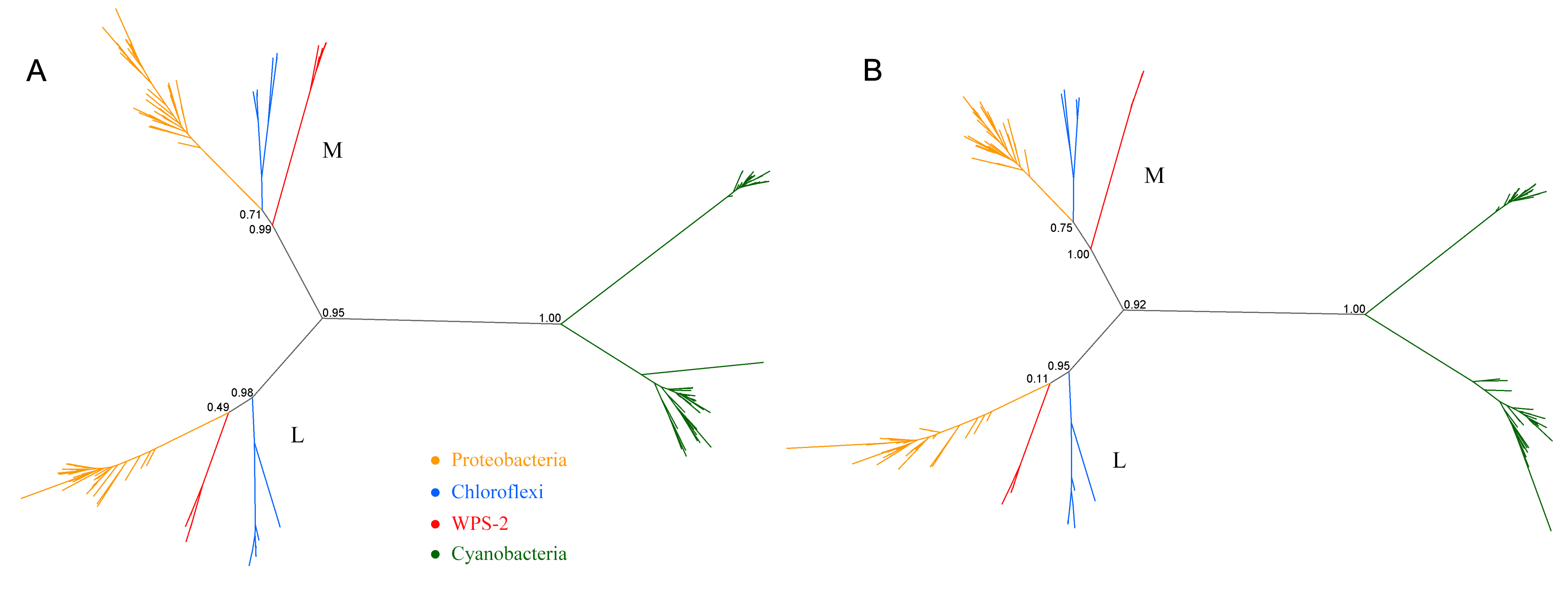
